# Disturbance opens recruitment sites for bacterial colonization in activated sludge

**DOI:** 10.1101/014456

**Authors:** David C. Vuono, Junko Munakata-Marr, John R. Spear, Jörg E. Drewes

## Abstract

Little is known about the role of immigration in shaping bacterial communities or the factors that may dictate success or failure of colonization by bacteria from regional species pools. To address these knowledge gaps, the influence of bacterial colonization into an ecosystem (activated sludge bioreactor) was measured through a disturbance gradient (successive decreases in the parameter solids retention time) relative to stable operational conditions. Through a DNA sequencing approach, we show that the most abundant bacteria within the immigrant community have a greater probability of colonizing the receiving ecosystem, but mostly as low abundance community members. Only during the disturbance do some of these bacterial populations significantly increase in abundance beyond background levels and in few cases become dominant community members post-disturbance. Two mechanisms facilitate the enhanced enrichment of immigrant populations during disturbance: 1) the availability of resources left unconsumed by established species and 2) the increased availability of niche space for colonizers to establish and displace resident populations. Thus, as a disturbance decreases local diversity, recruitment sites become available to promote colonization. This work advances our understanding of microbial resource management and diversity maintenance in complex ecosystems.

## Introduction

Dispersal of bacteria from regional pools to form local species assemblages is facilitated by a variety of mechanisms, including atmospheric transport via transcontinental winds, hitchhiking on migrating animals, circulating ocean currents, subsurface hydrological transport, and by anthropogenic means (Gilbert et al., 2009, 2012; Grossart et al., 2010; Harding et al., 2011; de Rezende et al., 2013; Smith et al., 2013; Bräuer et al., 2014; Müller et al., 2014). However, environmental filters, biotic interactions, relative fitness differences and stochastic processes (or combinations thereof) within heterogeneous ecological landscapes are the primary factors that dictate which organisms are recruited into local environments, and those that are not (Chesson, 2000; Hubbell, 2001; Adler et al., 2007). At local scales, species compete for limited resources and thus respond to and affect both the abiotic (e.g., resource gradients) and biotic (e.g., competition and predation) environment (Tilman, 1982; Chase and Leibold, 2003). Diversity within local environments is then maintained by stabilizing niche differences and relative fitness differences among species, which forms the basis of co-existence theory (Chesson, 2000). Such interactions feed back to influence the strength of environmental and biotic filters, which pose as colonization barriers to immigrant organisms.

Although niche and neutral paradigms were developed from the macro-ecological perspective, such patterns are also pervasive in microbial ecology. While high dispersal rates and rapid horizontal gene transfer among bacteria and archaea have been argued to erode population structure (Doolittle, 2009; Doolittle and Zhaxybayeva, 2010) (e.g., through the acquisition of plasmids that encode for pathogenicity, antibiotic production and resistance), wild bacterial populations are now known to be non-clonal and structured into ecologically and socially cohesive populations (Hunt et al., 2008; Cordero et al., 2012; Shapiro et al., 2012; Polz et al., 2013). Thus, the congruency between ‘macro’ and ‘micro’ ecological theory can allow for predictions regarding the role of colonization into stable and unstable (i.e., disturbed) ecosystems to better understand altered states in ecosystems. Such studies will improve our understanding of human health and our microbiome; it may be possible to divert or reverse disease states in humans that are linked to differences in bacterial community structures (Ley et al., 2005; Antonopoulos et al., 2009; Faith et al., 2011; Lozupone et al., 2012; Ridaura et al., 2013).

Stable ecosystems act as filters that prevent colonization of immigrant organisms. This is due to a lack of available resources and niche space, which are consumed and occupied by established species in local environments that provide localized ‘ecosystem services’ (Tilman, 2004). Thus, ecosystems that experience food web transitions either from primary or secondary succession (i.e., disturbance) may be more susceptible to colonization by immigrant organisms. Many examples of immigration and invasion exist in macro ecology (see Shea and Chesson, 2002). However, surprisingly few studies have attempted to characterize the patterns of colonization in microbial communities (Jones and McMahon, 2009; Wells et al., 2014). Jones and McMahon (2009) found minimal effect of immigration by atmospheric deposition of bacteria into freshwater seepage lakes; the composition of abundant immigrant communities was significantly more variable and did not overlap with the composition of abundant lake bacterial communities, most likely due to dispersal limitation. Microbial communities, however, are also known to have persistent ‘seed banks’ (Caporaso et al., 2011), which may contribute to microbial diversity maintenance through the increase in abundance and resuscitation of rare and dormant organisms into disproportionately active populations when conditions become optimal (Jones and Lennon, 2010).

Here we investigate the roles of immigration and colonization in a purely microbial ecosystem without dispersal limitation, before and during a long-term disturbance event, as well as during the successional periods that follow the disturbance. We define colonization as an increase in the abundance of taxa in a receiving environment that are also present in the source environment. Culture-independent methods were used to survey bacterial communities in a full-scale activated sludge bioreactor (Vuono et al., 2013) and its associated influent domestic wastewater as the sole immigration source. The disturbance imposed on the activated sludge bioreactor was a step-wise reduction in the operational parameter solids retention time (SRT), which is defined as average time that microorganisms are contained within the bioreactor and is equal to specific biomass growth rate (see Vuono et al., 2015 for complete description of the SRT disturbance). Briefly, the bioreactor was operated at a 30-day SRT (stable operating conditions), biomass wasting rate was then gradually increased to lower the SRT to 12 days, then to 3 days (maximum disturbance), and finally the system was returned to stable operating conditions at a 30-day SRT. All SRTs during the experimental period were operated for at least 4.3-times the SRT to allow for biomass acclimation. We hypothesized that colonization limitation would be greatest during stable operating conditions (i.e., before and after disturbance) and would decrease as the ecosystem progressed towards its maximum disturbance state. Such decreases in colonization limitation would thus be manifested as increases in taxonomic and phylogenetic similarity as well as an increase in the number of shared taxa between source (i.e., influent) and receiving (i.e., activated sludge) environments.

## Results

Sewage-derived (i.e., influent) bacterial communities were surveyed throughout the time-course in parallel with activated sludge communities that underwent disturbance and secondary succession (as reported by Vuono et al., 2015). A total of 135 samples were collected and the V4-V5 hypervariable region of the bacterial small-subunit (SSU) rRNA gene was sequenced from these samples. Of the 135 samples, 54 samples were collected from the influent and 81 samples were collected from the activated sludge bioreactor (triplicate samples per time-point), with 18 and 27 time-points over the 313-day time series, respectively. A total of 610,311 high quality amplicon reads were obtained (singletons removed) with sequence statistics of 1,384/13,904/4,258/4,521/2,187 (min/max/median/mean/SD, respectively). In the influent sequence library, 231,832 high quality sequences were recovered at an average read depth of 4,293 (±2,786 standard deviation or S.D.) sequences/sample and clustered into 1,395 OTUs of 97% sequence similarity. In the activated sludge sequence library, 378,479 high quality sequences were recovered at an average read depth of 4,672 (±1,499 S.D.) sequences per sample and clustered into 2,343 OTUs of 97% sequence similarity.

### Bacterial composition and diversity of sewage-derived and sludge communities

The majority of sequences from the influent libraries were mapped to bacterial phyla Firmicutes (62.0%), Proteobacteria (Alpha-2.0%; Beta-16.6%; Gamma-4.5%) and Bacteroidetes (9.5%) (Figure 1A top). The majority of Firmicute tag-sequences were associated with order/*family* Clostridiales (51.8%)/*Lachnospiracea* (26.9%), *Ruminococcaceae* (13.5%) and *Veillonellaceae* (6.6%) and Lactobacillales (7.4%) / *Carnobacteriaceae* (2.9%) / and *Streptococcaceae* (3.8%). Members of the phylum Bacteroidetes were mainly comprised of populations of order/*family* Bacteroidales (7.7%) / *Bacteroidaceae* (1.6%) and *Prevotellaceae* (5.3%) (Figure 1A). For the influent libraries, minor fluctuations in community composition were observed at the phylum and class levels and showed no significant differences in compositional similarity through time (Bray-Curtis R^2^_ADONIS_ = 0.16, *P* = 0.16; Morisita-Horn R^2^_ADONIS_ = 0.16, *P* = 0.248) or phylogenetic similarity (Weighted UniFrac, R^2^_ADONIS_ = 0.15, *P* = 0.25). These results indicate temporal stability of the source communities. Community composition of the activated sludge however, was greatly affected by the SRT treatments (as described by Vuono et al., 2015) and shifted such that community members within phyla Acidobacteria, Chloroflexi, Nitrospira, and Planctomycetes were present at high abundance under pre- and post-disturbance conditions (30-day SRTs) but were mostly washed out during the 3-day SRT (Figure 1A bottom). In contrast, organisms within bacterial clades Burkholderiales and Sphingobacteriales were opportunistic and became dominant members of the community during the 3-day SRT and biomass recovery periods (Vuono et al., 2015).

**Figure 1.**
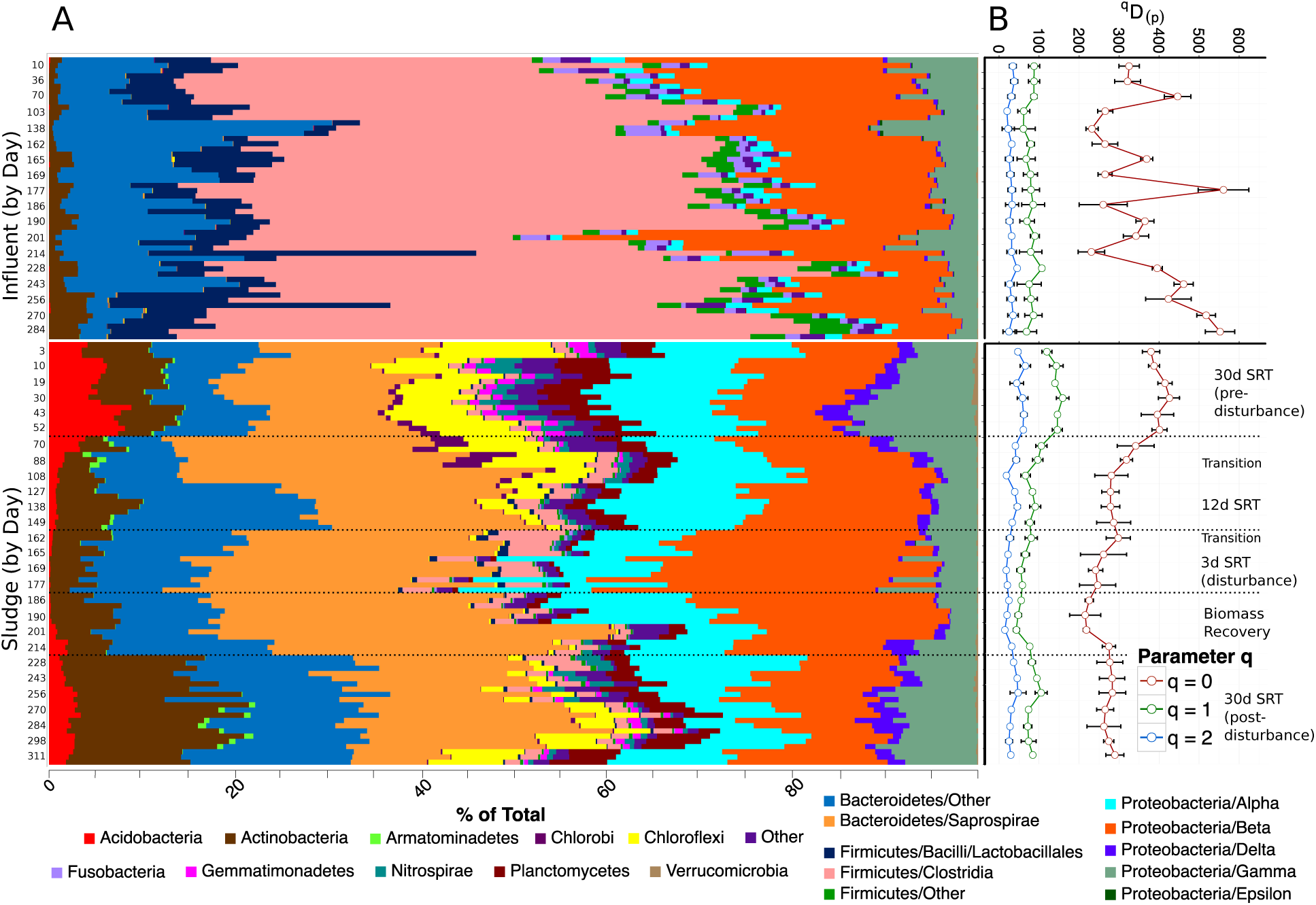
(A) Taxonomic summary of influent and sludge libraries, sorted by day of study with (B) corresponding Hill diversities. Error bars in B represent the 95% confidence interval estimated from a t-distribution. Sludge Hill diversities in B have been adapted from Vuono et al. (2015).

Mean Hill diversity of orders *^0^D*, *^1^D* and *^2^D*(*^q^D*_(**p**)_ for q=0,1,2) for the influent library were 366.6 (±109.4 SD), 79.5±19.9, and 30.4±10.4, respectively. Considerable variation in taxonomic richness, *^0^D*, was observed temporally compared to orders of diversity q=1,2 (Figure 1B), which indicates the lack of certainty in estimating species richness from sample data alone (Haegeman et al., 2013). First-order bacterial diversity, *^1^D*, of the influent communities was significantly lower than that of the activated sludge communities, pre- and post-disturbance (145.3±12.8 and 86.5±12.6, respectively as reported by Vuono et al. 2015) (Figure 1B). Diversity of order *^2^D*, which disproportionally emphasizes dominant species (i.e., a measurement of community evenness), was also lower in the influent than the activated sludge communities pre- and post-disturbance (59.9±9.0 and 37.7±10.3, respectively; Vuono et al., 2015). However, during the maximum disturbance state of the bioreactor (3-day SRT), diversity of order *^1^D* and *^2^D* in the activated sludge community was 21.4% and 23.0% lower than the influent diversities (62.5±6.0 and 23.4±2.8), respectively (Figure 1B).

### Disturbance drives phylogenetic community structure

To provide a phylogenetic context for the drastically different diversities across sample types (influent and activated sludge) and within SRT treatments (30 days to 12 days, to 3 days, followed by a recovery period and return to 30 days), the standardized effect size of each community’s mean phylogenetic distance (MPDSES) was calculated (Table S1). MPD_SES_ is a measure of phylogenetic relatedness that can be used to investigate contemporary ecological processes that may structure community composition (Webb, 2000), with positive and negative values indicating phylogenetic dispersion and clustering, respectively. Influent mean MPD_SES_ values calculated using the taxa.labels (presence/absence) and phylogeny.pool algorithms showed significant phylogenetic clustering throughout the time series (negative values). Similarly, MPD_SES_ values from the taxa.labels abundance weighted and independent swap algorithms were also consistently negative but did not suggest significant clustering at all time points throughout the time-series (Table S1). Based on the taxa.labels algorithm with presence/absence option, mean MPD_SES_ values of the activated sludge increased significantly as the disturbance progressed: 0.89±0.62 (pre-disturbance) to a maximum of 3.23±0.48 during the 3-day SRT and recovery period followed by a return to pre-disturbance levels, 1.50±0.80. The 3-day SRT MPD_SES_ values describe significant phylogenetic dispersion (i.e., limiting similarity) (MPD_SES_=2.95±0.57, *p*<0.005) and indicate that co-occurring species are significantly less related than expected by chance. These results are supported by a parallel increase in the probability of interspecific encounter (PIE), the probability that an organism of one species would encounter an organism of a different species (defined as 1-Simpson concentration; Hurlbert, 1971), during the 3-day SRT (0.017±0.003 to 0.041±0.007). Conversely, pre- and post-disturbance communities were randomly spread across the phylogenetic tree (MPD_SES_ values approaching zero), which indicates randomly assembled communities relative to the three null models tested. However, during the 3-day SRT (Table S1), a transition from random dispersion to significant overdispersion was detected for each of the null models, with the exception of the independent swap algorithm. The inability of the independent swap algorithm to detect niche-based community assembly is likely a result of its known conservatism (Kembel, 2009; Armitage et al., 2012), but does not change our interpretation of phylogenetic structure.

Phylogenetic clustering and overdispersion are commonly interpreted as the result of environmental (i.e., habitat filtering) and biotic (i.e., competition) filters (Webb, 2000). However, we use caution in naming any particular mechanism that describes phylogenetic community structure because several processes (e.g., phenotypic attraction and convergent evolution versus niche conservatism of functional traits) may produce similar tree topologies such as phylogenetic overdispersion. Thus, caution is warranted when interpreting the mechanisms of community assembly based on phylogenetic community structure, particularly in the absence of trait information. This is due to the difficulty in distinguishing the various combinations of environmental filters acting on organisms, stabilizing niche differences and the relative fitness differences among bacterial species (HilleRisLambers et al., 2012).

### Activated sludge communities become more similar to influent during disturbance

Bacterial communities within the bioreactor during each SRT treatment stage were compared to the influent (global comparison) to elucidate their taxonomic and phylogenetic similarity. We hypothesized that taxonomic and phylogenetic similarity would be greatest during the 3-day SRT because during this period resource availability is greatest and concentration of organisms in the sludge is lowest (Vuono et al., 2015), which lends to increased colonization probability of arrival organisms (Tilman 1994, 2004). Under all SRT treatments, sludge communities were significantly different from their immigration source (Table 1). However, for all metrics, the value of the R^2^_ADONIS_ statistic decreased to its lowest value during the 3-day SRT, which indicates a moderate increase in similarity (i.e., decreased variance between treatment types by sequential sum of squares). To quantify this similarity, the taxonomic and phylogenetic distance between the sludge and influent communities over time is plotted in Figure 2. The mean phylogenetic distance (Weighted UniFrac) between influent and bioreactor communities (0.63±0.06) was not significantly lower during the 3-day SRT (blue box Figure 2), with pre-and post-disturbance distances of 0.64±0.03 and 0.66±0.04, respectively. These results indicate that overall phylogenetic similarity of abundant organisms between influent and sludge did not increase during the 3 day SRT. Conversely, when rare taxa were weighted equally (unweighted UniFrac), the mean phylogenetic distance of influent and sludge communities decreased (0.83±0.01 pre-disturbance to 0.68±0.02 during disturbance). The same trend was observed with metrics of compositional similarity (Bray-Curtis and Morisita-Horn), which began to decrease during the 12-day SRT sludge wasting conditions (days 70 of study period) and decreased significantly more during the 3-day SRT. These results indicate an increase in similarity between the influent and bioreactor communities during the disturbance.

**Table 1.** Permutational MANOVA (ADONIS) analysis of taxonomic (Bray-Curtis and Morisita-Horn) and phylogenetic (Weighted UniFrac) similarity between source (i.e., influent) communities and their respective sludge communities, by SRT treatment. Significance in the ADONIS R^2^ statistic is based on 999 randomizations.

| Comparison | Bray-Curtis<br>$R^2$ | Morisita-Horn<br>$R^2$ | Weighted UniFrac<br>$R^2$ |
| --- | --- | --- | --- |
| 30 Day SRT vs. Influent | 0.553 | 0.697 | 0.718 |
| 12 Day SRT vs. Influent | 0.493 | 0.619 | 0.695 |
| 3 Day SRT vs. Influent | 0.384 | 0.512 | 0.615 |
| Biomass Recovery vs. Influent | 0.456 | 0.584 | 0.653 |
| 30 Day SRT (Recovery) vs. Influent | 0.551 | 0.682 | 0.715 |

**Figure 2.**
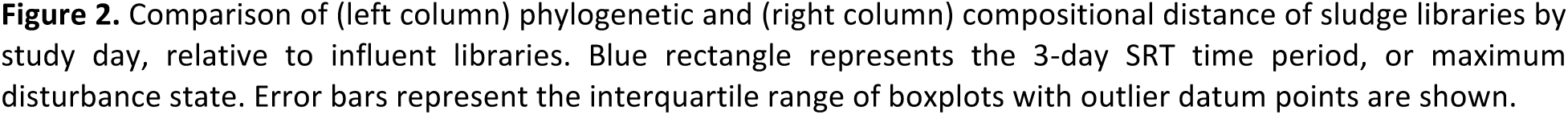
Comparison of (left column) phylogenetic and (right column) compositional distance of sludge libraries by study day, relative to influent libraries. Blue rectangle represents the 3-day SRT time period, or maximum disturbance state. Error bars represent the interquartile range of boxplots with outlier datum points are shown.

Colonization of individual taxa increased with disturbance progression. Overall, 40.1% of the OTUs were shared between influent and all sludge treatments and transition periods (Figure S1). However, because phylogenetic and compositional metrics (Figure 2) both showed small increases in similarity, it was unclear which contributed more to the increase in similarity: abundant or rare organisms. To answer this question, multivariate cutoff level analysis (MultiCoLA) was used to quantify the contribution of rare taxa to taxonomy-based beta-diversity metrics. Briefly, MultiCoLA removes the least abundant taxa at increasing proportions and correlates each reduced dataset to the full dataset to determine how each successive omission of rare taxa influences beta-diversity. This analysis revealed that each reduced dataset strongly correlated with the full dataset with the removal of up to 50% of the least abundant OTUs (Figure S2).

With this information, the dataset could be quantitatively reduced (hereby referred to as the reduced dataset) to more easily visualize the contribution of abundant and rare influent OTUs that may be shared with the sludge. A two-dimensional hierarchical clustering combined with a heatmap of OTU occurrence frequencies was plotted (Figure S3) to reveal which OTUs had similar co-occurrence patterns between sample groups. The 100 most abundant OTUs were sub-sampled to visualize the dynamics of individual taxa and the relative impacts of colonization during the disturbance (Figure 3). Hierarchical clustering of samples revealed that sample types were split between influent and sludge. Sludge samples clustered based on their respective SRT operation and the biomass recovery period, while influent samples did not cluster based on any particular structure (i.e., samples were distributed randomly in a single cluster). The lack of organized structure within the influent cluster supports a consistent influent community composition throughout the study period. The distribution of OTUs was also split between two main clusters. One cluster contained 42 OTUs that were consistently found at high abundance in the influent but low abundance in the sludge (lower left and right quadrants, respectively; Figure 3). The second cluster contained 58 OTUs that were either absent in the influent and abundant in the sludge or abundant in both libraries (upper left and right quadrants, respectively; Figure 3). The clustering patterns within this cluster were mainly driven by the OTU occurrence frequencies in the sludge, in response to disturbance and recovery (below), which are representative of ecological strategies as defined by Evans and Wallenstein (2014).

**Figure 3.**
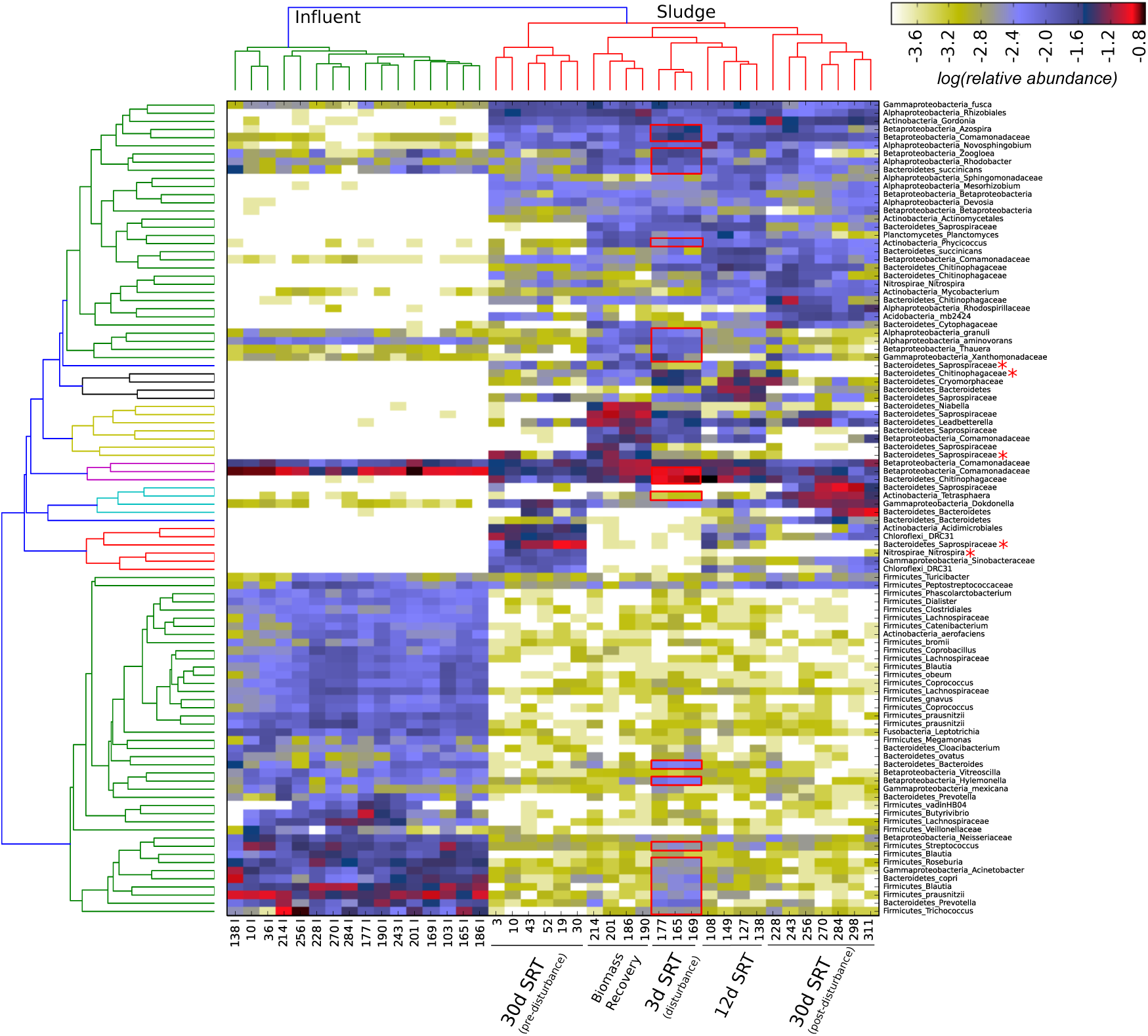
Heatmap and cluster analysis (Bray-Curtis; complete linkage) of the 100 most abundant OTUs in influent and sludge libraries; relative abundances are log transformed for visualization purposes. OTUs (y-axis) and samples (x-axis) are clustered based on similar abundance and occurrence patterns. Taxonomic identification of each OTU is shown on the right sidebar, starting with phylum (or class for Proteobacteria) and ending with the highest taxonomic classification assigned for that OTU. Red boxes capture the influent OTUs that increased in abundance in the bioreactor during the 3-day SRT, although other groups increased during 12-day SRT and biomass recovery as well. Red asterisks highlight the OTUs that did not recolonize to a comparable amount after disturbance based on limited representation in the sludge.

Several patterns of OTU occurrence frequencies are evident in the heatmap. These include (1) OTUs consistently present in significant proportions in influent, sludge, or both (generalists); (2) sludge OTUs that significantly decreased during the disturbance followed by recovery to higher levels; (3) loss of some organisms after disturbance (red asterisks); (4) addition of some organisms after disturbance; (5) lack of dependence of some OTUs on SRT. Of the sludge OTUs that were lost after the disturbance, it is possible that these organisms did not recolonize because no sequences from these OTUs were detectable in the influent. However, it is difficult to know if lack of recolonization was solely due to absence of a particular OTU in the influent because it is impossible to detect all organisms in any given sample.

If influent organisms were to establish as the activated sludge community members, the signature of successful colonization could be identified as a group of OTUs that were present in the influent, also present in the sludge, but increase in abundance beyond background levels (30-day SRT) only during the 3-day SRT (maximum disturbance). Potential colonizers would also have to be consistently abundant beyond background levels throughout the 3-day SRT. This suggests that potential colonizers have overcome a decay rate of 0.2 d^-1^, which is derived from activated sludge model 2 (ASM2d) based on oxygen uptake under starvation (Henze et al., 1999). Of the 100 most abundant OTUs, 23 were identified to have increased significantly in abundance above 30-day SRT levels (red boxes Figure 3). Thirty additional OTUs were also identified to have increased in abundance during the 3-day SRT, above 30-day SRT levels. In addition, all of the OTUs that were identified as having these abundance patterns (53 in total) were significantly enriched beyond background levels, as determined by a two-part statistic for comparison of sequence variant counts (*p*-values < 0.01; Figure 4). This collection of OTUs, albeit small, explains our findings from the distance-based taxonomic (Bray-Curtis, Morisita-Horn) and unweighted phylogenetic (UniFrac) calculations; combinations of abundant and rare influent taxa contribute to the increase in similarity during the disturbance periods, particularly during the 3-day SRT. However, it is still unclear why in the case of the weighted phylogenetic metric, 30-day SRT communities did not differ substantially from the 3-day SRT communities; it is possible that the phylogenetic composition of the abundant OTUs overwhelmed this signal.

**Figure 4.**
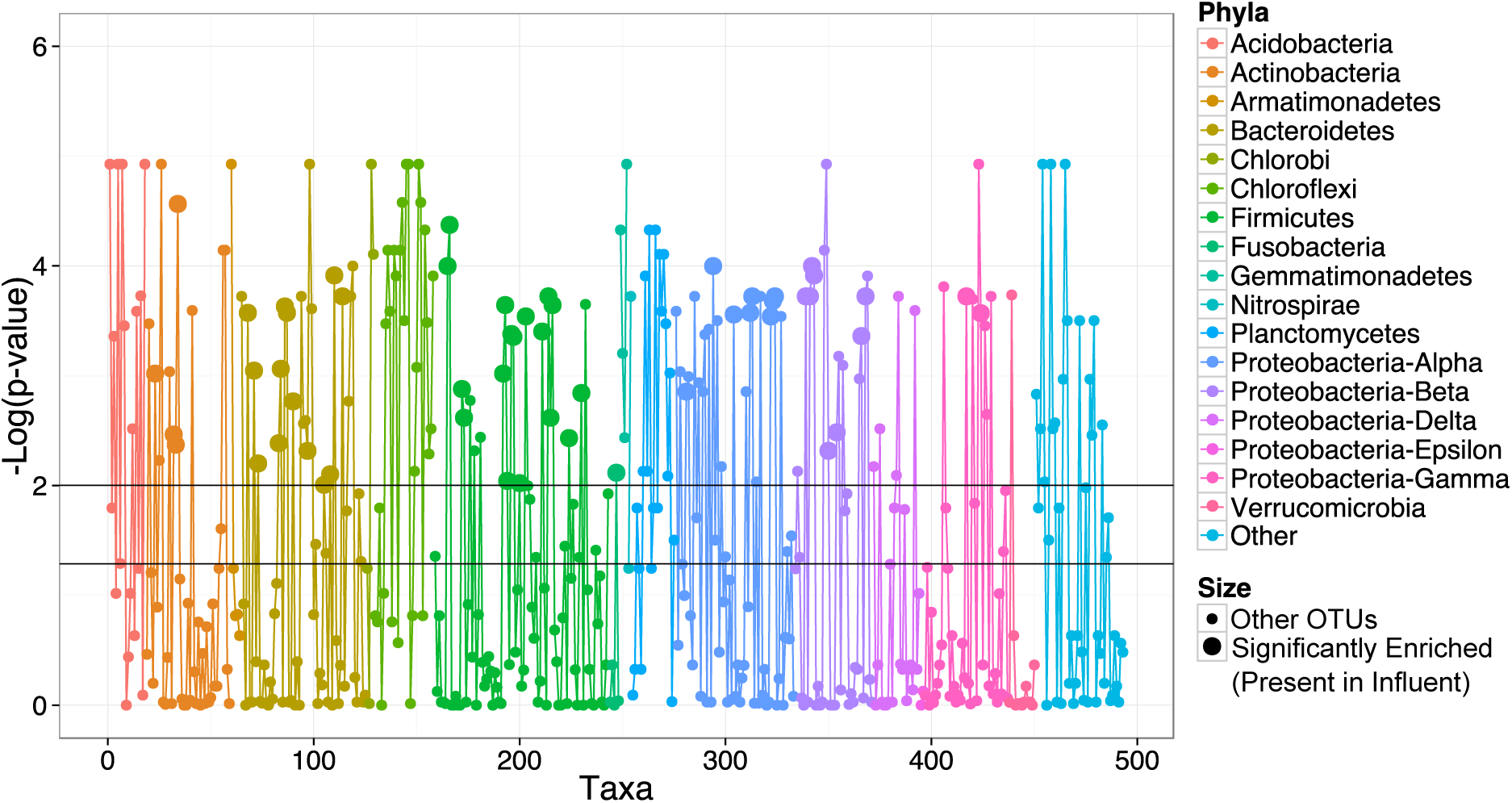
Manhattan plot, which displays negative log transformed p-values from all two-part tests for each OTU detected between 30-day and 3-day SRT operations. Values above 2 (top horizontal line; α = 0.01) represent OTUs that have significantly different abundance distributions, either enriched or depleted during the 3-day SRT (bottom horizontal line represents α = 0.05). The distinction between enriched or depleted OTUs is shown in Figure S4. Fifty-three OTUs (Table S2) were identified as being significantly enriched in the bioreactor from the influent during the 3-day SRT (represented as large data points). Colors correspond to different phyla and are ordered by taxonomy line. Phyla in the “Other” category (<1% abundance) are comprised of Spirochaetes, Synergistetes, Cyanobacteria, Elusimicrobia, Fibrobacteres, and candidate divisions BHI80-139, BRC1, FBP, GN02, GN04, GOUTA4, MVP-21, NKB19, OD1, OP3, SR1, TM6, TM7, WPS-2, and WS3.

### Network-based analysis reveals higher OTU co-occurrences between disturbance and influent communities

To visualize the degree of connectedness between samples as a function of shared OTUs, a network was built of the most prevalent OTUs (~2% or 198 OTUs), which had ≥150 sequences detected study-wide, including replicates (Figure 5). In the network, samples that cluster more closely together have more OTUs shared between them. The network clearly shows that samples cluster into two main groups, either influent or sludge (denoted by edge color radiated from sample nodes, with green and blue lines representing sludge and influent samples, respectively). All of the influent samples cluster closely together with little deviation between them. Conversely, as the disturbance progresses, the 3-day SRT sludge samples deviate from the 30-day SRT cluster and approach the influent cluster. This shift in position indicates that the 3-day SRT sludge samples share more OTUs with the influent than do the 30-day SRT treatments. Each sample node radiates edges to OTUs (outer nodes) that are either shared or not shared, independent of treatment. 65% of the OTUs in this analysis were shared between influent and sludge. The bacterial phylum that shared the most OTUs between sample types was Firmicutes (55 OTUs in total) with the remaining OTUs mapped to Proteobacteria (42 OTUs in total, mostly within the Alpha, Beta and Gammaproteobacteria classes), Bacteroidetes (22 OTUs in total) and Actinobacteria (8 OTUs in total). The OTUs most frequently shared between influent and sludge, and containing the largest edge-weights (i.e., the average number of sequences in an OTU) were mapped to the cosmopolitan clade of *Comamonadaceae* (OTU 1 and 3, Figure 3 and Table S2). Other shared and prevalent OTUs between influent and sludge were *Faecalibacterium prausnitzii, Lachnospiracea/Blautia sp*. and *Prevotella copri* (4, 5, and 23 OTUs). In summary, network-based analysis reinforces our observations of increased similarity between influent and sludge during disturbance and provides a robust method to visualize this similarity based on shared OTUs between baseline operational conditions and maximum disturbance state.

**Figure 5.**
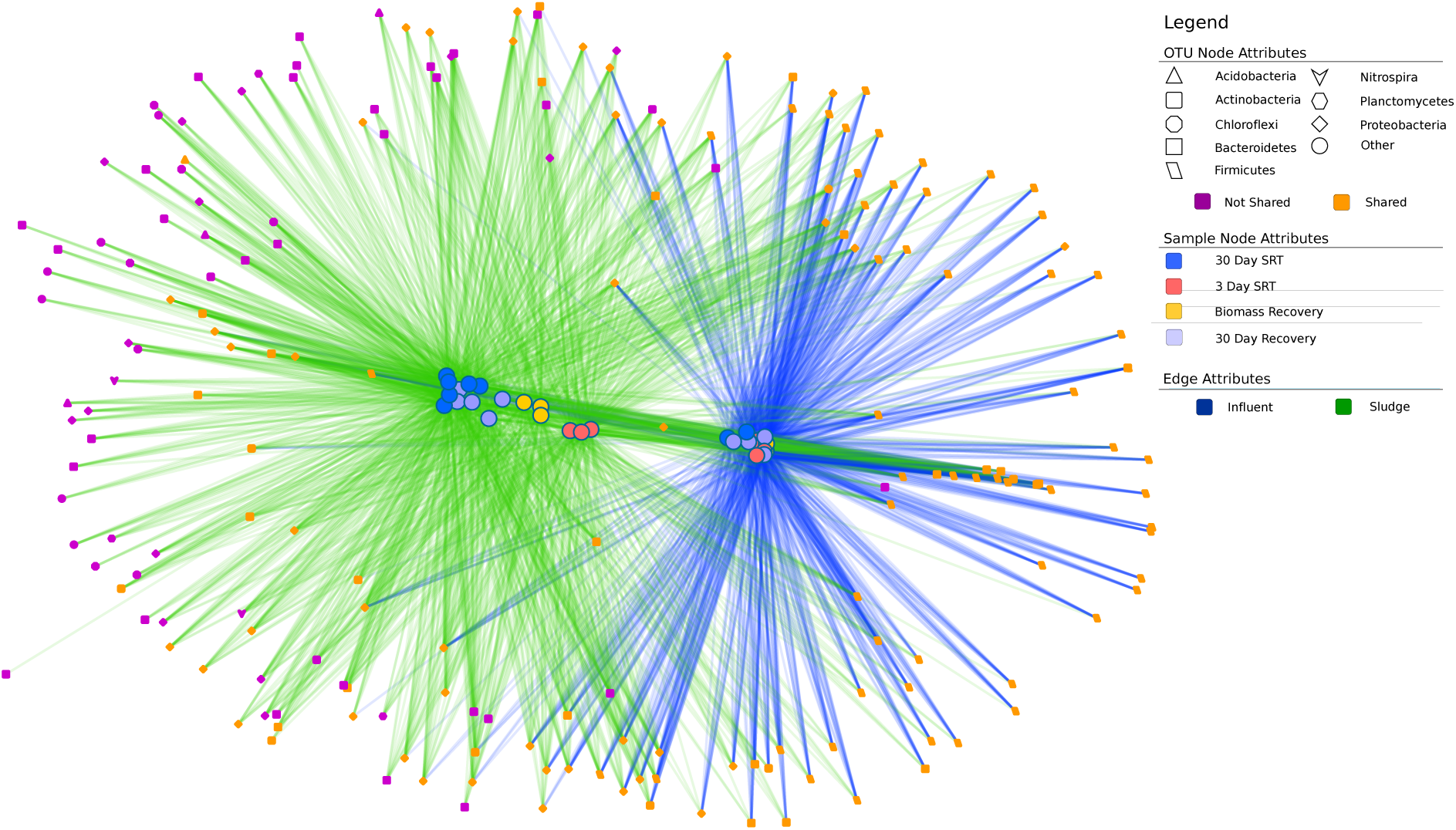
Shared OTU network analysis of the most abundant OTUs (capturing those influent OTUs that were consistently present throughout the study period) during the 30-day SRTs (pre- and post-disturbance; blue and light blue sample nodes, respectively), 3-day SRT (max-disturbance; red sample nodes) and biomass recovery period (orange sample nodes) (12-day SRT and transition periods have been removed for clarity). Sample nodes (inner) and OTU nodes (outer) are connected by edges, which indicate whether those OTUs are shared by Influent, Sludge, or both. Node shape indicates the phylum level classification of the OTU.

## Discussion

In this work, the influence of bacterial colonization into an ecosystem (activated sludge bioreactor) was measured through a disturbance gradient (successive decreases in solids retention time) relative to stable operating conditions. Results show an increased abundance of organisms in the sludge that were compositionally similar to those in the influent during the maximum disturbance state. Our results support the hypothesis that disturbance opens up recruitment sites for bacterial colonization. These results add to the predictive framework of microbial resource management within engineered microbial systems, but also provide insights for how to manage and promote healthy microbial communities (e.g., within the human microbiome, for individuals with diseased states associated with unstable community structures). These results also shed light on the similarities between ‘macro’ and ‘micro’ ecology, which show similar patterns of dispersal and colonization of ecosystems during disturbance (Tilman, 1994, 2004). In addition, the stability of the bioreactor ecosystem (as measured by the range of ecosystem processes and decrease in diversity; Vuono et al., 2015) was compromised and manifested as a decreased resistance to invasion (i.e., niche opportunity) (Shea and Chesson, 2002; Shade et al., 2012). During stable operating conditions (flanking treatments before and after disturbance), the opposite trend was observed: taxonomic and phylogenetic similarities between influent and sludge decreased as did the number of shared OTUs (both weighted abundance and presence/absence) between influent and sludge sequence libraries. These results could be interpreted under the ‘species-sorting’ metacommunity paradigm (Leibold et al., 2004), as suggested by Jones and McMahon (2009), where spatial niche partitioning is emphasized over dispersal dynamics.

Tilman (2004) describes such systems under the more parsimonious framework of stochastic niche theory, where invading species can survive on resources left unconsumed by established species. In the case of the bioreactor communities (i.e., sludge), established species were washed out of the system during the disturbance and thus dampened the effects of recruitment limitation. Conversely, recruitment of influent bacteria into the activated sludge communities was considerably lower during the pre- and post-disturbance (30-day SRTs). Several lines of evidence support these observations, based on the predictions of stochastic niche theory. First, pre- and post-disturbance Hill diversities (for both *^1^D* and *^2^D*) were considerably higher in pre- and post-disturbance than the maximum disturbance communities. Recruitment limitation is not a direct consequence of highly diverse communities, but rather a consequence of the low levels of available resources unconsumed by organisms within highly diverse communities (Tilman, 2004). Second, the concentrations of bacteria in the bioreactor during pre- and post-disturbance were an order of magnitude higher than the disturbance communities (Vuono et al., 2015), which indicates an environment highly saturated with microorganisms. In addition, the Food:Microorganism (F/M) ratio was an order of magnitude higher during the disturbance than during normal operating conditions (Vuono et al., 2015). These results all indicate increased substrate availability during the disturbance and corresponding higher proportion of colonization in the context of stochastic niche theory.

Within the context of metacommunity theory, an activated sludge bioreactor can be characterized as a source/sink system: source habitats (influent) are not constrained by finite growth and thus overflow into sink habitats (i.e., bioreactor) (Leibold et al., 2004), where sludge wasting rates control which microbes persist based on their growth rates. We see evidence for such dynamics with the depletion of several OTUs (Figure 3, red asterisks) that were washed out of the bioreactor system during the disturbance but were not repopulated after the disturbance due to their absence within the influent. For example, two functionally important nitrite-oxidizing *Nitrospira*, phylogenetically constrained to sublineages I and II (OTUs 63 and 85, respectively), were present and abundant (as high as 4% of the community) in the bioreactor before the disturbance (Vuono et al., 2015). Not surprisingly, neither *Nitrospira* OTU was detected in the influent since *Nitrospira* are obligate aerobes, not thriving in environments such as the human gut or urban sewer water infrastructure (Lücker et al., 2010; Vandewalle et al., 2012). However, the population of *Nitrospira* sublineage I was able to reestablish (i.e., regrow) after the disturbance because it was not completely washed out of the bioreactor. Yet, *Nitrospira* sublineage II was completely washed out of the system as early as the 12-day SRT and did not repopulate to a comparable amount in the bioreactor after the disturbance, most likely due to its absence in the influent.

Our results indicate that a high proportion of low-abundance OTUs were shared between influent and 30-day SRT communities (Figures 3 and S1B). This may be indicative of a taxa-time relationship, or an increase in observed taxa richness with increasing time of observation (Preston, 1960); the bioreactor was operated for an 8-month period under a 30-day SRT prior to the start of the experiment (Vuono et al., 2015), which may have allowed for more influent taxa to accumulate as rare community members in the bioreactor with time. The accumulation of influent organisms as low-abundance activated sludge members, which displayed opportunistic growth during the 3-day SRT, may be an example of how dormancy and rarity contribute to diversity maintenance within microbial ecosystems (Jones and Lennon, 2010). We cannot definitively say in the current study that low-abundance community members are indeed dormant using the current methods, as rare members may be active but may not contribute to overall community structure and function. Nonetheless, environmental (abiotic) and biotic filters (species interactions), which form the basis of co-existence theory, play a large role in regulating the colonization ability of immigrants to shape local species assemblages (Chesson, 2000).

Many microbial studies have aimed to determine the relative contributions of such ecological forcing mechanisms by calculating the phylogenetic relatedness of communities relative to null expectations (Horner-Devine and Bohannan, 2006; Armitage et al., 2012; Stegen et al., 2012; Wang et al., 2012; Meuser et al., 2013; Evans and Wallenstein, 2014). Phylogenies that are either clustered or overdispersed have traditionally been interpreted to reflect environmental filters (e.g., habitat filtering) or biological interactions (e.g., competitive exclusion), respectively, that are responsible for the assembly of local communities (Webb, 2000). However, given the assumption that phylogenetic relatedness is a proxy for trait and niche similarity, the declaration of assembly mechanisms should be used cautiously (Mayfield and Levine, 2010; HilleRisLambers et al., 2012). This may be especially true for bacteria, due to their high dispersal rates and genetic recombination rates (Hunt et al., 2008; Polz et al., 2013; Cordero and Polz, 2014). In the case of the sludge communities, we observed phylogenetic overdispersion during the maximum disturbance state (3-day SRT) followed by a shift to random dispersion during normal operating conditions (30-day SRT). Interestingly, Loreau and Mouquet (1999) found that immigration plays a large role in competitive communities, which would coincide with our observations if indeed the overdispersed community at 3-day SRT could be explained by competitive exclusion. Nonetheless, whatever the mechanism that caused this shift in phylogenetic structure, ecosystems processes in the bioreactor, such as nitrification, denitrification and phosphorus removal, were greatly impaired by the disturbance (Vuono et al., 2015). Thus, future studies should elucidate if phylogenetic overdispersion is a property of disturbed ecosystems, as a way to gauge ecosystem health and resistance to invasion.

## Experimental Procedures

### Bioreactor operation

A full-scale decentralized activated sludge bioreactor treating domestic wastewater was operated through time and at 3 different solids retention times (SRTs): 30, 12 and 3-days as described by Vuono et al. (2015). Briefly, hydraulic retention time (HRT: 24 hrs.) and SRT were decoupled through biomass separation (ultrafiltration membranes with a nominal pore size of 0.05μm) in a sequencing batch membrane bioreactor. SRT was managed through controlled wasting of return activated sludge.

### Sampling, Amplification, and Sequencing

All biological samples for each time-point observation were collected in triplicate. Activated sludge samples were collected as described by Vuono et al. (2015). Influent samples were collected from treatment plant headworks post primary screening. All samples were stored at -20 °C prior to processing. DNA was extracted within one month of sampling with MoBio’s PowerBiofilm DNA extraction kit (Carlsbad, CA) following manufacturer’s protocol with a 1-minute bead-beating step for cellular lysis. Barcoded small sub-unit (SSU) rRNA gene primers 515f-927r were incorporated with adapter sequences for the GSFLX-Titanium platform of the Roche 454 Pyrosequencing technology as described by Vuono et al. (2015).

### SSU rRNA processing, quality control, and diversity analysis

SSU rRNA gene amplicons generated from pyrosequencing were binned by barcode and quality filtered with the ‘split_libraries.py’ script in the Quantitative Insights Into Microbial Ecology (QIIME v1.8-dev) software (Caporaso et al., 2010). Sequences with errors in the barcode or primer, lengths shorter than 400nt or longer than 460nt, a quality score <50, homopolymer run greater than 6 nt, or which contained ambiguous base-calls were discarded from downstream analysis. The remaining sequences were corrected for homopolymer errors with Acacia (Bragg et al., 2012). Chimeric sequences were identified with UCHIME under reference mode and *de novo* mode (Edgar et al., 2011). The remaining 613,321 sequences were processed in Mothur (Schloss et al., 2009) as outlined by Schloss SOP version data 2014/15/02. Sequences were clustered into OTUs at 97% sequence similarity using average neighbor method. Taxonomic classification of de-replicated sequences was implemented with the RDP Classifier (Wang et al., 2007) at 80% bootstrap confidence against the Greengenes 16S rRNA database (13_5 release) (DeSantis et al., 2006). Singletons were discarded from downstream analysis prior to diversity calculations. A phylogenetic tree was constructed from the filtered alignment with FastTree. Unweighted and weighted UniFrac distance matrices were calculated from the phylogenetic tree. Beta diversity metrics (i.e., Bray-Curtis dissimilarities and Morisita-Horn distances) were derived from a rarefied operational taxonomic units (OTU) table from the sample with lowest sequencing depth. Diversity analyses were performed in the R environment for statistical computing, QIIME, Mothur, and Explicet (See *SI Materials and Methods* for complete description of diversity analysis). All sequence data generated in this study are publicly available in the MG-RAST database under the accession No. 4569429.3.

## Acknowledgments

The material presented is based in part upon work supported by the National Science Foundation under Cooperative Agreement EEC-1028968. We owe many thanks to Dave Armitage, William Navidi, Tzahi Cath, Terry Reid, Lloyd Johnson, Aaron Saunders, Dean Heil, and Chuck Pepe-Ranney for thoughtful discussions and suggestions on the manuscript.

## Conflict of Interest

The authors declare no conflict of interest.

